# Preclinical Evaluation of a Zotarolimus-Eluting Stent for Intracranial Application: A 30-Day GLP Canine Study

**DOI:** 10.1101/2025.11.25.690612

**Authors:** Ibrahim A. Bhatti, Fareed Jumah, Syed A. Gillani, Ravi Nunna, Inamullah Khan, Austin Rolfes, Bharathi Jagadeesan, Ameer E. Hassan, Adnan I. Qureshi, Farhan Siddiq

**Affiliations:** Department of Neurosurgery, University of Missouri, Columbia, MO; Zeenat Qureshi Stroke Institute and Department of Neurology, University of Missouri, Columbia, MO; Office of Medical Research, University of Missouri, Columbia, MO; Departments of Radiology, Neurology and Neurosurgery, University of Minnesota, MN; Department of Neurology, Valley Baptist Medical Center, Harlingen, Texas, US

**Keywords:** Intracranial Arteriosclerosis, Drug-Eluting Stents, Zotarolimus, Canine, Animal, Model, Safety

## Abstract

**Background:** Zotarolimus-Eluting Stents (ZES) are increasingly used off-label for intracranial atherosclerotic disease (ICAD). However, foundational, Good Laboratory Practice (GLP) compliant preclinical safety data specific to the cerebral vasculature is lacking. Therefore, we conducted the first systematic evaluation of the technical feasibility, anatomic features, and 30-day clinical safety profile of the Onyx Frontier™ ZES (Medtronic, Minneapolis, MN) in a GLP-compliant canine intracranial model.

**Methods:** In a prospective GLP study, a 2.0 × 8.0 mm Onyx Frontier™ ZES was implanted in the basilar artery of twelve healthy mongrel canines pre-treated with dual antiplatelet therapy. Vertebral artery tortuosity was assessed via angiography and graded as Type I (non-tortuous/straight), Type II (moderately tortuous/S-shaped), or Type III (severely tortuous/coiled). The primary endpoints were procedural success and 30-day safety, assessed via angiography, serial neurological examinations, and comprehensive clinical monitoring.

**Results:** Technical success was achieved in 100% of animals (12/12). The cohort exhibited challenging anatomy, with a mean basilar artery diameter of 1.39 ± 0.10 mm. Tortuosity analysis revealed that 33.3% of animals were Type I (mild), 58.3% were Type II (moderate), and 8.3% were Type III (severe). Post-procedure, mild vasospasm was observed in all cases, with only one canine requiring intra-arterial verapamil. A single case (8.3%) of self-limiting contrast extravasation occurred without clinical sequelae. At 30 days, there was no mortality, and no animals suffered a neurological adverse event. Minor clinical events were unrelated to the device, and comprehensive clinical pathology panels revealed no evidence of systemic toxicity or a significant inflammatory response.

**Conclusion:** This first-of-its-kind preclinical study demonstrates that implantation of the Onyx Frontier™ ZES is technically feasible with a 100% success rate in a sub-2mm vessel model with significant tortuosity and has an excellent 30-day safety profile with no clinical evidence of neurotoxicity. These findings provide foundational preclinical data supporting further investigation of this technology for treating ICAD in human subjects.

## Introduction

Intracranial atherosclerotic disease (ICAD) represents a significant cerebrovascular pathology responsible for approximately 8-10% of all ischemic strokes in North America and up to 46.6% of strokes and transient ischemic attacks in Asian populations.^1^ The American Heart Association/American Stroke Association (AHA/ASA)’s currently recommends aggressive medical management^2^ for ICAD, which the faces the challenge of a high risk of recurrent strokes in the stenosed vessel territory, which has been estimated to be 23% at one year,^3^ most of which are disabling or fatal.^4^ This has driven the evolution of endovascular therapies. However, early trials with bare-metal stents (BMS) were limited by high peri-procedural risks, in-stent restenosis (ISR), and showed non-superiority to medical management, as highlighted by the SAMMPRIS, VISSIT and CASSIS trials.^5^ ISR rates with BMS have been seen to be as high as 30%.^6^ Drug-eluting stents (DES), particularly Zotarolimus-eluting stents (ZES), are a more recent development in the endovascular treatment of ICAD. ZES have shown promise in coronary arteries by inhibiting neointimal hyperplasia, the primary driver of restenosis, and have become the standard of care in coronary vascular pathology.^7^ This has led to increasing off-label clinical use in the intracranial circulation.^7–11^ The Onyx Frontier™ (Medtronic, Minneapolis, MN) ZES is a newer-generation DES with features designed for enhanced deliverability in tortuous intracranial anatomy.^12^ Despite its growing off-label use for ICAD, systematic, Good Laboratory Practice (GLP)-compliant preclinical safety data, a mandated requirement from the U.S. Food and Drug Administration (FDA) prior to pivotal clinical trials,^13^ are lacking.

Previously, Sirolimus-eluting stents (SES) have been evaluated in preclinical studies and have shown an acceptable neurotoxicity and clinical safety profile.^14–16^ None of those reports provided extensive technical details or elaborated on the complete angiographic cerebrovascular anatomy of the preferred canine model for intracranial applications. Furthermore, no such studies exist evaluating ZES in a canine model. Recognizing this literature gap, the primary goal of this study was to generate this foundational data for technical feasibility and 30-day safety of the Onyx Frontier™ ZES in a GLP-compliant, translationally relevant canine basilar artery model. A specialized canine model was selected because it allows for assessment of device deliverability in a challenging, tortuous anatomy mimicking human neuro-navigation^17^. The study was designed to collect systematic short and long-term safety, toxicology and neurotoxicity data that cannot be obtained with this level of control in existing clinical studies. Complete neurotoxicity and detailed pharmacokinetics data will be reported elsewhere upon completion of the study. We have also described the angiographic cerebrovascular anatomy in this canine model for future application. The results of this preclinical study may be used as standardized regulatory support for a future Investigational Device Exemption (IDE) application in human clinical trials.^18,19^

## Methods

### Study Design and Regulatory Compliance

This investigation is a prospective, GLP study designed to evaluate the safety and technical success of the Onyx Frontier™ ZES in a canine intracranial model. The study was conducted in compliance with the United States Food and Drug Administration (FDA) GLP Regulations (21 CFR Part 58) under protocol OFZ001-IS05. All animal procedures were reviewed and approved by the Test Facility’s Institutional Animal Care and Use Committee (IACUC).

### Animals and Husbandry

Twelve healthy adult male mongrel canines (Canis lupus familiaris) were sourced from an approved vendor. The canine model was selected based on its well-documented suitability for neuroendovascular procedures, including a gyrencephalic brain, the absence of a rete mirabile, and vessel sizes that accommodate human-grade delivery systems.^17^ The canine basilar artery model was selected for its strong anatomical and procedural comparability to human cerebral vasculature. Its vessel caliber is comparable to that of human intracranial arteries.^17^ Furthermore, the canine cerebrovasculature is characterized by inherent tortuosity, particularly in the carotid and vertebral arteries, which replicates a primary navigational challenge in human neurointerventions.^17,20^ Successfully deploying a stent in this model therefore provides a robust and clinically relevant demonstration of the device’s deliverability and flexibility, directly addressing a critical hurdle in the endovascular treatment of ICAD.

Animals were group-housed under standard conditions in accordance with the Guide for the Care and Use of Laboratory Animals. Twelve mongrel canines (25-50 kg) were assigned to three cohorts (n=4 per cohort) with scheduled termination time points at 30-, 90- and 180-days post-implantation. The present report details the technical feasibility and 30-day safety outcomes from the entire cohort (N=12); data from the 90- and 180-day cohorts, including comprehensive histopathology and pharmacokinetic analysis, will be reported elsewhere.

### Anatomical and Technical Rationale for Implant Approach

The implant strategy was informed by consulting prior literature regarding the anatomy of the canine cerebral vasculature.^17,20^ The posterior circulation is the dominant source of flow to the canine Circle of Willis, with robust posterior communicating arteries (PCoA) ensuring bilateral filling of the anterior circulation upon basilar artery injection.^17^ Furthermore, the cervical internal carotid artery (ICA) is notably tortuous and prone to catheter-induced vasospasm, making consistent and safe intracranial access via an anterior approach tough.^17^ In contrast, the vertebrobasilar system, while also tortuous, provides a more feasible and stable conduit for device delivery. The distal basilar artery was selected as the target vessel for this study since anatomy of other large intracranial arteries in this model is not feasible for endovascular application.

The vertebral arteries in the dog arise from the subclavian arteries, ascend through the cervical transverse foramina, and, together with the ventral spinal artery contribute to a spinal arterial circle in the vertebral canal at the level of the atlanto-occipital articulation. The rostral portion of this spinal arterial circle extends a short distance into the cranial cavity and continues as the basilar artery (BA).^21,22^ The BA in the dog gives off cerebellar and brainstem branches that correspond functionally and topographically to human posterior inferior cerebellar (PICA-type), anterior inferior cerebellar (AICA-type), and superior cerebellar (SCA-type) arteries, and frequently issues medullary and pontine perforators and several accessory cerebellar branches. Canine specimens commonly show variations in exact origins (e.g., PICA arising from either BA or a VA), and significant extracranial–intracranial anastomoses (occipital–vertebral and orbital/anastomotic channels) are repeatedly reported in the anatomy literature.^21,22^ We have described this typical canine posterior circulation anatomy in **Figure** 1, a digital subtraction angiogram taken from one of our canines. The vertebral arteries often follow a tortuous course, providing a rigorous model for testing the navigability of endovascular devices.^17^

**Figure 1:**
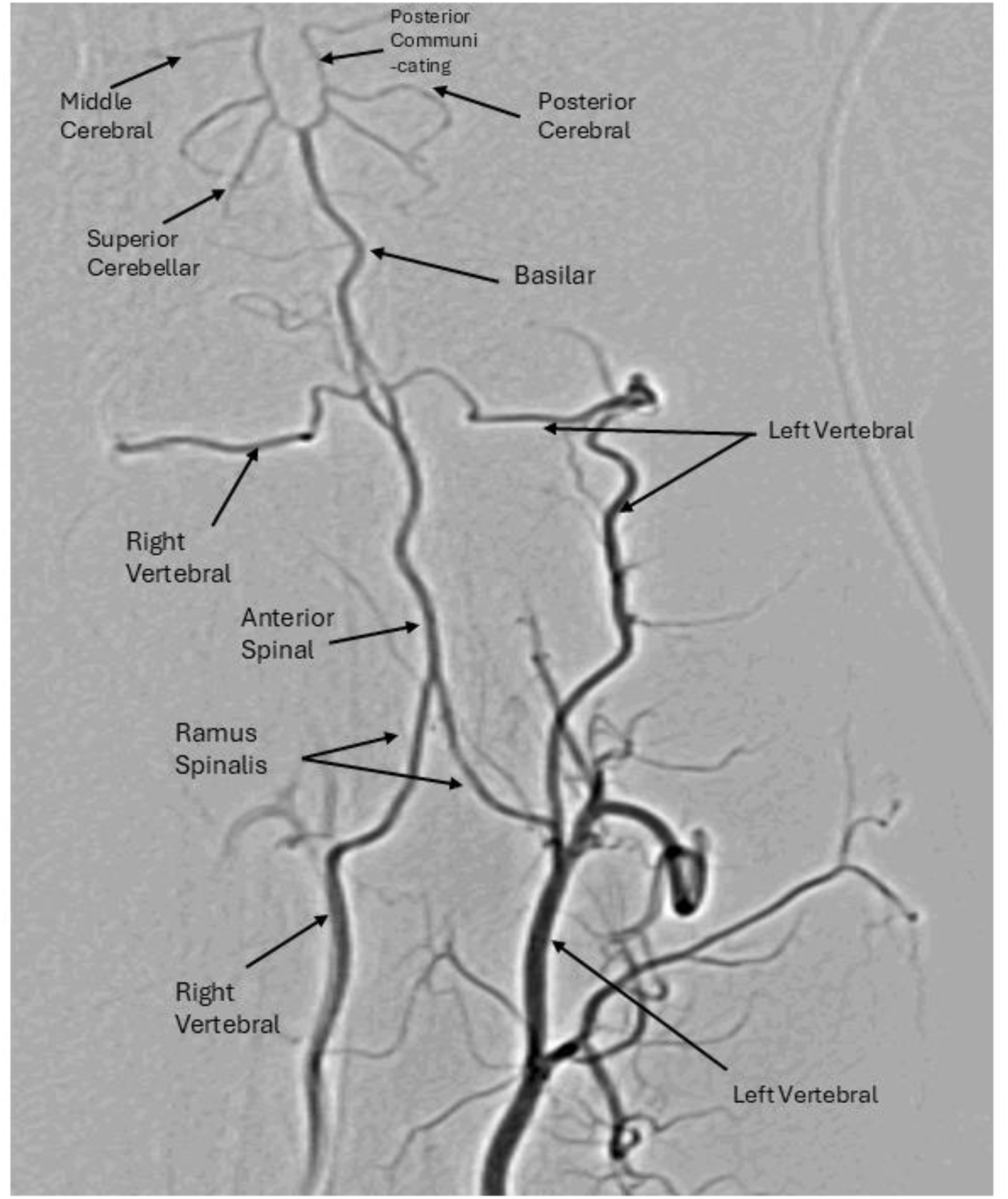
Labeled reference digital subtraction angiogram (DSA) of the canine posterior circulation. The image demonstrates the typical vertebrobasilar anatomy, including the vertebral artery (VA), basilar artery (BA), superior cerebellar artery (SCA), and posterior cerebral artery (PCA).

### Test Article

The study utilized the Onyx Frontier™ DES, a ZES (2.0 mm diameter, 8.0 mm length) with a drug concentration of approximately 1.6 µg/mm²

### Rationale for using 2 x 8 mm design

The 2.0 x 8 mm Onyx Frontier™ ZES was selected for its low-profile design, which is engineered for enhanced trackability in tortuous anatomy.^23^ The Onyx Frontier™ ZES features a modern thin-strut design with a thickness of 81 µm, fabricated from a cobalt-based alloy.^24^ While comparable in strut thickness to the Wingspan (75 µm),^25^ the stent displays a substantial reduction in strut thickness compared to older-generation drug-eluting stents which utilized stainless steel struts ranging from 132 to 140 µm (e.g., Cypher, TAXUS).^26^

An important safety consideration for intracranial stenting is the mitigation of ostial obstruction to critical perforating arteries. Anatomically, the intracerebral perforators are divided into proximal and distal segments.^27^ The larger lateral group of lenticulostriate arteries exhibits mean diameters of 510 μm for the proximal segment and 470 μm for the distal segment, while the smaller medial group averages between 260–280 μm.^27^ Consequently, the 81 μm strut thickness of the Onyx Frontier™ ZES is roughly 3-to-6 times smaller than the average luminal diameter of these critical perforators, significantly minimizing the risk of flow obstruction into these vital side branches. This favorable design profile, which ensures the strut is substantially smaller than even the smallest major perforators (≥80 μm), supports the safe application of this stent in the challenging sub-2mm vasculature characteristic of intracranial atherosclerotic disease, as has already been demonstrated in human subjects.^28^

### Pre-Medication and Anesthesia

After a minimum 5-day pre-treatment with dual antiplatelet therapy (DAPT; clopidogrel 75 mg and aspirin 81 mg PO SID), animals received pre-medication on the day of the procedure. This included nifedipine (30 mg PO) for antispasmodic prophylaxis, cefpodoxime (5-10 mg/kg PO) as an antimicrobial, and carprofen (2.5-5.0 mg/kg PO) for analgesia and anti-inflammatory effects.

General anesthesia was induced with midazolam (0.1-0.2 mg/kg IM) / butorphanol (0.05-0.1 mg/kg IM) and propofol (2.0-8.0 mg/kg IV to effect) and maintained with isoflurane (0-5% in 100% O₂).

### Technique of Endovascular Stent Implantation Procedure

Under general anesthesia and sterile technique, surgical cutdown was performed for right femoral artery access. Using the modified Seldinger technique, a 5 Fr femoral sheath was placed. Under fluoroscopic guidance, a 5 Fr MPD guide catheter was advanced into the dominant vertebral artery over a 0.035 Glidewire®. After baseline cerebral angiography and quantitative vessel analysis (QVA) of the basilar artery, the ZES stent delivery system was advanced over a micro-guidewire. The stent was deployed in the basilar artery by slow balloon inflation. Immediate post-implant angiography confirmed accurate deployment and full stent expansion. A second angiogram at 3-5 minutes confirmed patency and the absence of in-stent thrombosis. Finally, the femoral artery was ligated, and the access site was surgically closed.

### Vertebral Artery Anatomy, Tortuosity Assessment, and Access Strategy

Pre-procedural angiography was used to assess two key anatomical variables: (1) the pattern of vertebral artery dominance, classified as right-dominant, left-dominant, or co-dominant (balanced); and (2) the degree of cervical vertebral artery tortuosity. Vascular tortuosity was graded using the vertebral artery tortuosity classification described by Ryu et al.^29^ This classification, developed for posterior-circulation interventions, defines three morphological types as, Type I) Nontortuous: straight or gently C-shaped curvature without acute angulation (< 90°), Type II) Moderately tortuous: S-shaped configuration or a single site of acute angulation (< 90°), and Type III ) Severely tortuous: coiled or kinked vessel, or acute angulation at > 1 location. Each canine vertebral artery was assessed on anteroposterior and lateral DSA projections from the ostium to the intracranial V4 segment. Two neurointerventionists independently assigned grades; discrepancies were resolved by consensus. The femoral access site and guiding catheter trajectory were selected based on this anatomical assessment. A standardized approach was employed where the right vertebral artery was accessed preferentially in all cases of right-dominance and co-dominance. The left vertebral artery was selected for access only in cases of clear left-dominance.

### Post-Operative Care and Follow-up

Post-operative analgesia was provided with buprenorphine and tramadol. The dual antiplatelet regimen was continued throughout the study duration. Prophylactic antimicrobials and non-steroidal anti-inflammatory drugs were administered for 5 and 3 days, respectively. Humane endpoints were pre-defined in the study protocol, including weight loss exceeding 20%, severe anemia (hematocrit <15%), or unremitting neurological deficits/pain unresponsive to analgesia. Animals were monitored daily, with formal neurological examinations performed daily for the first three days and weekly thereafter.

### Specimen Collection and Analysis

Blood samples were collected at baseline, and on days 1, 3, 7, 14, and 30 for pharmacokinetic analysis of zotarolimus. Standard hematology and serum chemistry panels were performed at baseline, day 14, and termination. C-reactive protein (CRP) was assessed at baseline and on days 3, 7, and 14.

### Termination and Histopathology

At the scheduled endpoints, animals will be euthanized under deep anesthesia via intravenous potassium chloride injection. The brain and basilar artery will be perfusion-fixed. The stented vessel segment will be processed for plastic embedding, sectioned (proximal, mid, distal), and stained with Hematoxylin and Eosin (H&E) and Movat’s Pentachrome. Brain tissue will be processed for standard paraffin histology. Since the scope of this report is technical feasibility, detailed angiographic anatomy of canine model and 30-day clinical results, pharmacokinetics and histopathology results, which are not available at this time, may be reported elsewhere upon completion.

## Results

### Technical Feasibility and Angiographic Outcomes

Stent implantation was technically successful in all 12 canines (100%). Individual anatomical characteristics and procedural details are summarized in **Table 1**. The mean basilar artery diameter was 1.39 ± 0.10 mm (range 1.25–1.56 mm), confirming deployment in a sub-2 mm vessel environment. The cohort exhibited a spectrum of vertebral artery tortuosity: 4 animals (33.3%) had mild (Type I) tortuosity, seven animals (58.3%) had moderate (Type II) tortuosity, and one animal (8.3%) had severe (Type III) tortuosity. Vertebral artery dominance was right or co-dominant in 10 animals (83.3%) and left-dominant in 2 animals (16.7%). In accordance with the access protocol, right femoral access was used in 10 cases and left femoral access in the 2 cases with left-dominant anatomy. The procedure was successful even in animals with challenging anatomy, including moderate and severe vertebral artery tortuosity.

**Table 1:** Individual Animal Anatomical and Procedural Details (N=12)

| **Animal ID** | **Basilar Artery Diameter (mm)** | **Vertebral Artery Dominance** | **Vertebral Artery Tortuosity Type** | **Access Side** |
| --- | --- | --- | --- | --- |
| 25C0128 | 1.30 | Co-dominant | I | Right |
| 25C0129 | 1.34 | Co-dominant | III | Right |
| 25C0131 | 1.56 | Co-dominant | II | Right |
| 25C0133 | 1.25 | Left Dominant | I | Left |
| 25C0134 | 1.42 | Co-dominant | I | Right |
| 25C0135 | 1.47 | Co-dominant | II | Right |
| 25C0136 | 1.26 | Right Dominant | I | Right |
| 25C0154 | 1.30 | Co-dominant | II | Right |
| 25C0155 | 1.41 | Left Dominant | II | Left |
| 25C0164 | 1.36 | Co-dominant | II | Right |
| 25C0165 | 1.54 | Co-dominant | II | Right |
| 25C0166 | 1.45 | Co-dominant | II | Right |

### Vertebral Artery Tortuosity

Post-deployment control angiography demonstrated successful stent placement with excellent wall apposition in all cases. No perforator or branch vessel occlusion was noticed. Immediate brisk anterograde flow was confirmed through the stented segment. There were no instances of acute in-stent thrombosis, vessel dissection, or perforation. **Figure 2** demonstrates pre- and post-stenting angiograms for two animals from the cohort, demonstrating the technical success of the stenting and the tortuous canine posterior circulation, highlighting the model’s procedural complexity.

**Figure 2:**
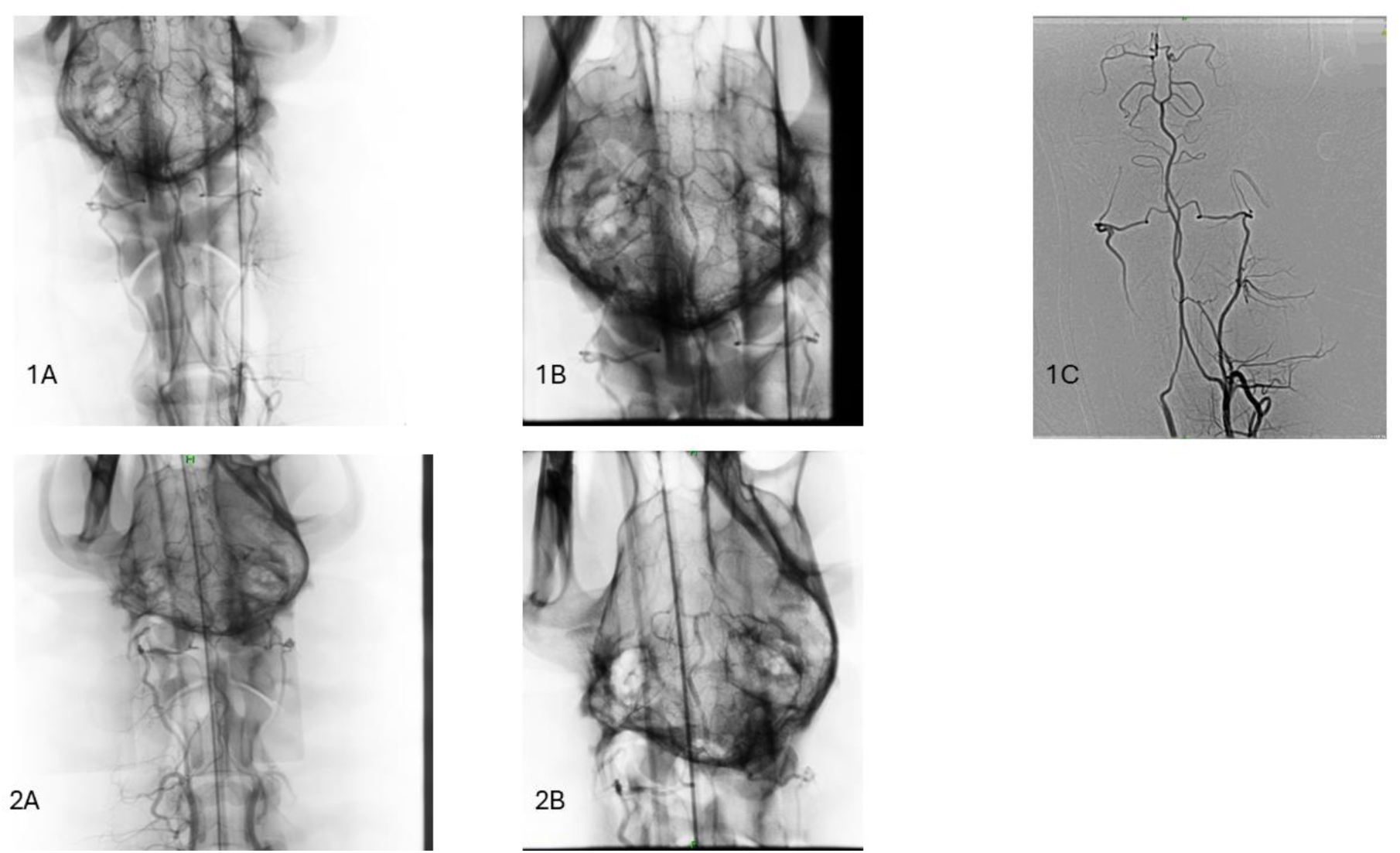
Composite of representative angiographic results demonstrating technical success in challenging anatomy. **1A-1C:** showing implantation in an animal with Type I (mild) tortuosity. **1A)** pre-stenting angiogram of animal ‘1’. **1B)** post-stenting angiogram of animal ‘1’. **1C)** Digital subtraction angiogram of animal ‘1’. **2A-2C:** showing implantation in an animal with Type II (moderate) tortuosity. **2A)** pre-stenting angiogram of animal ‘2’. **2B**) Post-stenting angiogram of animal ‘2’.

Procedure-related complications: One procedural complication was noted: a single case (8.3%) of suspected minor contrast extravasation (animal 25C0133), which was observed during the procedure and deemed not clinically significant, requiring no further action. Follow-up angiography did not show any dissection or vessel injury. Mild, transient vasospasm was noted post-deployment in some animals; only one was treated with 5mg of intra-arterial verapamil. All the remaining ones resolved spontaneously in 3-5 minutes.

### 30-day Clinical and Neurological Safety Outcomes

All 12 animals remained in good general health at the 30-day endpoint. The clinically significant observations noted during the first 30 days were infrequent, minor, and determined to be unrelated to the implanted device or the endovascular procedure. Animal 25C0129 (30-day cohort): Experienced a recurrence of a pre-existing pododermatitis on Day 11, which resolved by Day 24 with topical treatment. Animal 25C0165 (30-day cohort): A transient heart murmur was noted under anesthesia on Day 0, which resolved spontaneously. On Day 1, a post-operative subconjunctival hemorrhage was observed, which resolved without intervention by Day 16. Animal 25C0128 (180-day cohort): Sustained minor wounds and transient inappetence from an altercation with a cage mate on Day 24, which were resolving with standard veterinary care. Animal 25C0133 (180-day cohort): A suspected contrast extravasation in the basilar artery was noted intra-operatively, which required no further action and resulted in no clinical sequelae. No other clinically significant abnormalities or adverse neurological signs were observed in any of the twelve study animals.

All animals remained neurologically intact throughout the study period, with no deficits noted on serial examinations. The minor, non-neurological clinical events observed were successfully managed and considered unrelated to the implanted device. All animals maintained or gained weight, indicating good systemic health. A summary of the 30-day cohort outcomes is provided in **Table 2**. The mean weight change for this cohort was 3.18% ± 2.18% (range 0.3% to 5.6%), indicating good systemic health.”

**Table 2:** Summary of 30-Day Cohort Clinical Outcomes (N=4)

| **Animal ID** | **Neurological Status** | **Clinically Significant Observations (Unrelated to Device)** | **Outcome** | **Weight Change (%)** |
| --- | --- | --- | --- | --- |
| 25C0129 | Intact | Pre-existing pododermatitis | Resolved with topical treatment | +3.4% |
| 25C0165 | Intact | Transient heart murmur;<br>Subconjunctival hemorrhage | Resolved spontaneously | +0.3% |
| 25C0166 | Intact | None | - | +5.6% |
| 25C0164 | Intact | None | - | +3.4% |
|  |  |  |  | **Mean:**<br>3.18% ±<br>2.18% |

### Clinical Pathology

Comprehensive clinical pathology panels were analyzed for all animals. No clinically significant trends indicative of systemic toxicity was observed. A summary of the findings is presented in **Table 3**.

**Table 3:** Summary of Clinical Pathology Findings (All Animals, 30-Day Period)

| **System** | **Key Parameters** | **Findings** |
| --- | --- | --- |
| Hematology | WBC, RBC, Platelets, Hemoglobin | All parameters remained within normal limits. No evidence of anemia, leukopenia, or thrombocytopenia. |
| Serum Chemistry | Liver Enzymes (ALT, ALP), Renal Markers (BUN, Creatinine), Electrolytes | Isolated, mild, and transient excursions noted (e.g., elevated ALP), consistent with normal biological variability. No evidence of systemic organ toxicity. |
| Inflammatory Markers | C-Reactive Protein (CRP) | No consistent trend observed across the cohort, indicating an absence of a significant, sustained systemic inflammatory response. |

## Discussion

This study provides the first GLP-compliant, preclinical evaluation of the Onyx Frontier™ ZES in a canine intracranial model. The primary findings, 100% technical success and an excellent 30-day safety profile, are critical, as they provide the foundational biological data to support the device’s widespread clinical use in ICAD. The technical success rate in the canine model, wherein the brain anatomy and cerebral vessel size are comparable to humans,^17,30^ is particularly meaningful given the anatomical challenges presented by vertebral artery tortuosity of the model and the sub-2 mm target vessel caliber. This corresponds to the success already seen in recent interventional studies in human intracranial vessels, including sub-2mm vasculature.^9,23,31^

The canine vertebrobasilar system is a well-established model, but it is not without procedural difficulties.^17^ The significant tortuosity and propensity for catheter-related vasospasm inherent to the canine neurovasculature have been described, as have the challenges with more rigid microcatheter systems, noting significant vessel straightening and even instances of subarachnoid hemorrhage attributed to catheter stiffness.^17^ In this context, the ability to successfully navigate and deploy the Onyx Frontier™ stent in all 12 animals without significant complication provides strong preclinical evidence for the flexibility and deliverability of the delivery system, highlighting its suitability for complex human intracranial anatomy. Care was taken to enroll large mongrels weighing at least 25kg. This was to ensure that the experimental animals had vessels of large enough caliber to accommodate endovascular intervention.^32^.

For similar reasons, in a prior large-scale preclinical investigation by Sun et al.,^16^ SES were studies in a large-sized canine model. Their study, which selected 154 Labrador Retrievers weighing over 25 kg, confirms the selection of large, adult canines as the gold standard for such evaluations, ensuring well-developed vasculature that closely approximates human intracranial vessel dimensions and tortuosity.^17^ Furthermore, the robustness of the canine model for replicating complex neurointerventional scenarios has been validated by studies such as that of Zhang et al.,^33^ who successfully developed a novel vertebral artery occlusion technique, underscoring the model’s capacity for precise endovascular manipulation.^33^

The high technical success observed in our challenging canine model may be attributed to the design characteristics of the current-generation Onyx Frontier™ ZES. While similar to its predecessor (Resolute Onyx™) in stent platform, polymer, and drug, the Onyx Frontier™ features design modifications such as increased catheter flexibility and a smaller crossing profile.^23^ These advancements, including a dual-layer balloon, are reported to provide a 16% improvement in deliverability over the previous platform without compromising radial strength.^34^ Our preclinical findings mirror the high success rate of the ZES platform in clinical studies. For example, a 2024 multicenter study by Chahine et al.^23^ demonstrated 100% technical success in 23 patients with acute ischemic stroke. Furthermore, the predecessor device, the Resolute Onyx™, has also shown high technical success rates (95%-100%) in clinical studies for symptomatic ICAD.^35,36^

Zotarolimus, the antiproliferative agent on the ZES, is a sirolimus analog optimized for stent-based delivery with a more rapid tissue uptake and shorter half-life, potentially reducing systemic exposure.^37–40^ The mechanistic action of zotarolimus, like sirolimus, involves mTORC1 inhibition to suppress neointimal hyperplasia.^37^ While neurotoxicity remains a theoretical concern for ‘limus’ drugs in the cerebral vasculature,^38^ our 30-day safety results (showing no clinical neurological complications) align with prior preclinical evidence from same family drugs. Levy et al.,^14,15^ and Sun et al.^16^ reported neurological safety of SES, noting that drug levels in brain tissue remained below concerning thresholds. It must be stated, however, that local drug concentrations in the adjacent brain tissue were not assessed as part of this 30-day report, which limits definitive conclusions on local neurotoxicity. Zotarolimus does possess a favorable pharmacokinetic profile with faster tissue uptake and a shorter half-life than sirolimus, potentially mitigating systemic exposure risks.^39^

The context for this preclinical work is defined by the severe limitations of current ICAD management. The results of the SAMMPRIS and VISSIT clinical trials showed high peri-procedural and recurrent stroke risk when using stenting with the older BMS devices, compared to the medical management groups,^41^ thus shaping the AHA/ASA guidelines for ICAD to recommend medical management and advice against endovascular treatment.^42^ However, medical management still carries a high recurrent stroke risk, with the SAMMPRIS medical arm showing a 15% primary endpoint rate and a 34.3% two-year re-stroke risk for patients with prior stroke.^43^ This unmet need has driven interest in DES. Clinical data, though off-label, has been highly promising. A randomized trial, the NOVA trial^44^ utilizing SES, showed lower recurrent stroke (0.8% vs. 6.9%) and ISR (9.5% vs. 30.2%) for DES over BMS. ZES literature is similarly favorable: a propensity-matched analyses showed ZES 1-year endpoints (11.5%) were significantly lower than the SAMMPRIS medical arm (28.1%). A 3-year propensity-matched comparison showed ZES (11.3%) outperformed both medical management (27.0%) and percutaneous transluminal angioplasty and stenting (27.8%) arms from SAMMPRIS.^40^ Despite this growing body of supportive clinical evidence, these devices are not FDA-approved for this indication. Therefore, our GLP preclinical safety study is intended to support further controlled investigation toward regulatory evaluation.

Several limitations of this study warrant consideration. First, the use of a healthy canine model, while essential for controlled safety assessment, does not replicate the human pathophysiology of ICAD. The absence of atherosclerotic plaque, calcification, and associated hemodynamic alterations limits direct extrapolation of these results to the treatment of diseased human arteries. Second, the sample size, though reasonable for an initial feasibility and safety study, provides limited power to detect uncommon adverse events, however, the cost of the GLP study, and availability of canines for any such applications are major limiting factors. Third, the 30-day follow-up period for this report is insufficient to evaluate long-term outcomes such as delayed restenosis or late stent thrombosis. However, the scope of this report was to demonstrate technical feasibility if new generation ZES strengthened by early clinical safety. Long term safety and toxicology results will be reported elsewhere. Finally, while no neurological or systemic toxicity was observed, drug concentrations in the adjacent brain tissue were not quantified as part of this 30-day analysis, precluding definitive conclusions on local neurotoxicity. The comprehensive histopathological and pharmacokinetic data from the 90- and 180-day cohorts, which will address these long-term endpoints, are pending and will be studied in future.

## Conclusion

This first-of-its-kind, GLP-compliant study demonstrates that intracranial implantation of the Onyx Frontier™ ZES is technically feasible, with a 100% success rate in a challenging tortuous cerebrovascular anatomy of large-animal model. The device is safe to deploy in sub-2 mm vessel diameter. The device and drug are associated with an excellent 30-day safety profile and no evidence of neurological adverse events or systemic toxicity. These findings provide essential preclinical safety data to support further controlled investigation toward regulatory evaluation of this technology for ICAD.

## Acknowledgments

None

